# Re-evaluating of Chemotaxis in *Acanthamoeba castellanii* and *Acanthamoeba polyphaga*: A Modern Perspective Using Advanced Imaging and Tracking Technologies

**DOI:** 10.1101/2025.10.29.683042

**Authors:** Viktor Hermaraj, Brendan W. Wren, Fauzy Nasher

## Abstract

Chemotaxis, the directed movement of an organism towards nutrients or away from noxious agents is a fundamental process for the survival of many microorganisms. We combined high-resolution imaging, microfluidic gradients, and frame-by-frame tracking to re-evaluate Acanthamoeba chemotaxis to microbial glycans (Mannan, Mannose, *N*-acetyl-d-glucosamine (GlcNAc), *N*-acetyl-muramic acid (MurNAc)) and peptides (*N*-formyl methionyl-leucyl-phenylalanine (fMLP) and Boc-Phe-Leu-Phe-Leu-Phe (BOC_FLFLF)). Our quantitative tracking results on *A. castellanii* confirm the core patterns in the original studies reported by Schuster and Levandowsky; attraction to fMLP and GlcNAc and lack of response to MurNAc or the peptide antagonist BOC-FLFLF, while revealing previously missed attraction to mannan. In contrast, *A. polyphaga* demonstrated a more restricted response, with significant chemotaxis observed only toward fMLP, and lack of motility in the presence of MurNAc or BOC-FLFLF. Notably, formyl peptide responses were differentially modulated: BOC-FLFLF reduced fMLP-induced directionality in *A. castellanii* without impairing motility, while in *A. polyphaga*, it suppressed both velocity and orientation. These distinct chemoattractant “signatures” are consistent with micro-niche adaptation, and we hypothesise that fine-scale tuning of receptor thresholds to local prey spectra contributes to the observed species differences rather than large-scale habitat segregation. By integrating advanced methodologies with classical paradigms, this study offers new perspectives on Acanthamoeba chemotaxis and supports emerging models of protist pattern recognition paralleling innate immunity.

## Introduction

*Acanthamoeba* species, including *A. castellanii* and *A. polyphaga*, are ubiquitous free-living protists that inhabit soil, water bodies, air, and human-made environments like swimming pools and plumbing systems (Rayamajhee et al., 2022). These organisms play vital ecological roles as bacterivores, shaping microbial communities and nutrient cycles (Mungroo et al., 2021; Price Christopher et al., 2024). They can be opportunistic pathogens capable of causing amoebic keratitis and granulomatous amoebic encephalitis, particularly in immunocompromised individuals or contact-lens wearers (Marciano-Cabral and Cabral, 2003; Zhang et al., 2023).

Pioneering research by Schuster & Levandowsky (1996) pioneered the quantitative study of *A. castellanii* chemotaxis, demonstrating directed migration toward stimuli such as *N*-formyl-methionyl-leucyl-phenylalanine, Lipopolysaccharides (LPS), lipid A, lipoteichoic acid, cyclic AMP, and *N*-acetylglucosamine (GlcNAc), but not toward Mannose, Mannosylated BSA, or *N*-Acetylmuramic acid (MurNAc) and Boc-Phe-Leu-Phe-Leu-Phe (BOC-FLFLF) (Schuster and Levandowsky, 1996).

Subsequent investigations revealed the presence of high-affinity mannose-binding proteins (MBPs) on *A. castellanii*, which facilitate adhesion to mannose-rich ligands on host cells and microbes, enhancing phagocytosis and cytotoxicity (Garate et al., 2006). Genomic and protein-level analyses have characterized varying mannose- and laminin-binding protein repertoires (MBP, MBP1, LBP) across different Acanthamoeba species and genotypes, suggesting evolutionary diversification of prey recognition systems (Clarke et al., 2013; Corsaro, 2022; Matthey-Doret et al., 2022). The *A. castellanii* genome also encodes additional candidate recognition proteins, including D-galactoside/L-rhamnose-binding lectins (RBLs), peptidoglycan recognition proteins (PGRPs), and H-type lectin domain proteins (Clarke et al., 2013; Nasher and Wren, 2024).

Recent work indicates that *A. castellanii* detects conserved microbe-associated molecular patterns (MAMPs), and that glycan presentation often governs recognition. Here we use “microbial glycans” to mean surface-exposed carbohydrate motifs on microbes, LPS inner cores/O-antigens, peptidoglycan (MurNAc–GlcNAc), capsular polysaccharides, β-1,3-glucans, and mannans/high-mannose N-glycans, that function as conserved MAMPs. For LPS, *E. coli* engineered to display only the Kdo◻–lipid A inner core were internalized at near wild-type levels, and preincubation with purified Kdo◻–lipid A, but not lipid A alone, blocked uptake, indicating that the two Kdo residues linked to lipid A are necessary and sufficient for recognition in this system.(Liu and Koudelka, 2023). Second, *A. castellanii* binds β-1,3-glucan on fungal cell walls, and masking these polymers diminishes uptake of Histoplasma (Ferreira Md et al., 2024). Third, uptake of *Campylobacter jejuni* depends on *O*-linked glycans on the major flagellin (FlaA); mutating the glycosylation sites abrogates this interaction (Nasher and Wren, 2023). Together, these reports point to a common logic: Acanthamoeba recognizes glycan-decorated microbial surface assemblies—LPS inner cores, glucan-rich walls, and *O*-glycosylated flagella—where the conserved appendage is the scaffold and glycan presentation fines tunes engagement (cell-cell recognition?). This pattern is consistent with a modular recognition system (lectin-like receptors for sugars alongside other sensors for peptides), paralleling PRR-based detection in multicellular hosts (Li and Wu, 2021; Nasher and Wren, 2024). At the molecular level, Acanthamoeba mannose-binding proteins and mammalian lectins (e.g., mannose receptor, collectins) show domain-level similarities and functional analogies, consistent with convergent evolution of microbial recognition pathways (Ferreira et al., 2022). Understanding chemotaxis in free-living amoebae matters ecologically, these protists structure bacterial communities, and evolutionarily, because their MAMP-guided navigation parallels PRR-based discrimination in innate immunity. By quantifying ligand-specific guidance in two species, we connect functional foraging strategies to conserved recognition logics that predate animals (Nasher and Wren, 2024).

Despite technological advances tracking microbes, comparative chemotaxis assays across *Acanthamoeba* species remain scarce. Prior studies have largely focused on *A. castellanii* (Schuster and Levandowsky, 1996; Kuburich et al., 2016), and quantitative chemotaxis of *A. polyphaga* towards microbial glycans has not been reported, even though lectin-mediated glycan recognition has been described for this species (Elloway et al., 2004). Here, we provide the first rigorous, quantitative comparison of *A. polyphaga* chemotaxis to this ligand panel under microfluidic gradients, using defined microbe-associated ligands and modern micro-fluidic imaging with frame-by-frame tracking. We use “chemotaxis” to include both attraction and chemo-repulsion; however, because a canonical chemorepellent for *Acanthamoeba* under microfluidic gradients has not been established, our assays focus on attraction metrics. We revisit classical chemotaxis assays using time-lapse imaging and quantitative cell tracking to analyse *A. castellanii* responses to mannan, mannose, GlcNAc, MurNAc, fMLP, and the antagonist BOC-FLFLF. Importantly, we also conduct a direct comparative chemotactic profiling of *A. polyphaga*, offering fresh insights into species-level sugar sensing and lectin-mediated navigation.

Our comprehensive chemotaxis metrics, covering multiple microbial-associated ligands and peptide stimuli, provide a foundational framework for understanding prey discrimination by *Acanthamoeba* in environmental and pathogenic contexts. By applying modern live-tracking technology to classic assays, this work paves the way for exploring how MAMP-driven microbial recognition in protists parallels the evolution of innate immune discrimination in warm-blooded hosts.

## Methods

### Slide Preparation and Media Composition

All experiments were conducted using Ibidi μ-Slide Chemotaxis chambers (Ibidi, Germany), which are specifically designed for live-cell imaging and chemotactic assays. Peptone-Yeast-Glucose (PYG) medium was freshly prepared and syringe-filtered (0.22 µm) for amoebal growth. For chemotaxis assays, glucose-free defined medium (ADM: NaCl: 1 g, KCl: 0.04 g, MgSO◻ · 7H◻O: 0.02 g, CaCl◻: 0.01 g, Na◻HPO◻: 0.14 g, KH◻PO◻: 0.06 g, FeCl◻: 0.5 mg at pH 7.5), which comprises only defined salts and iron. The omission of glucose and other nutrients ensured that observed amoeboid movement reflects ligand-driven migration rather than background metabolic stimuli.

### Amoeba Culture and Preparation

*Acanthamoeba castellanii* CCAP 1501/10 and *Acanthamoeba polyphaga* CCAP 1501/14 *◻Culture collection of Algae and protozoa (Scottish Marine Institute)]* were cultured axenically in PYG medium at 25◻°C under standard atmospheric conditions in 75◻cm^2^ Corning™ tissue culture flasks containing 25◻mL medium. Initial cultures were incubated for four days to reach log-phase growth. Amoebae were harvested at a concentration of approximately 2.5 × 10◻ cells/mL, as determined by manual counting using a haemocytometer.

Prior to each experimental session, cultures were gently dislodged from flask surfaces by tapping the flasks against the workbench. The cell suspension were centrifuged at 4000 ×◻g for 2◻min. The supernatant was discarded, and the amoeba pellet was resuspended in 25◻mL ADM. Cell dispersal was facilitated by pipetting and vortex mixing to ensure homogeneous distribution.

### Slide Inoculation and Chemotactic Assay Setup

Ibidi μ-Slides were loaded using the manufacturer’s protocol. Briefly, the central channel was inoculated with amoeba suspensions of either *A. castellanii* or *A. polyphaga*. One chamber reservoir well always contained ADM alone, while the right well was loaded with ADM supplemented with test stimulus at 1◻mM concentration. Stimuli used include Mannan, Mannose, *N*-acetylglucosamine (GlcNAc), *N*-Acetylmuramic acid (MurNAc), *N*-formyl-methionyl-leucyl-phenylalanine (fMLP), and the antagonist Boc-Phe-Leu-Phe-Leu-Phe (BOC-FLFLF). Mannan, due to its higher potency, was used at 10–20◻nM.

### Live-Cell Imaging

Live-cell imaging was performed using a Zeiss confocal microscope (LSM880) controlled via ZEN software. Time-lapse recordings were acquired over 2500 frames at a rate of one frame every 2.5◻s (total duration ∼104◻min) at objective 10x. Raw imaging data were exported from ZEN as uncompressed AVI files for subsequent analysis.

### Image Processing and Cell Tracking

Time-lapse image analysis was conducted using Fiji (a distribution of ImageJ) with pre-installed plugins. For quantitative tracking, a Substack was generated with the input range set from frames 1–2000 at 25-frame intervals (i.e., every 62.5◻s). This yielded an 80-frame AVI suitable for chemotaxis analysis. If amoeba motility was delayed, Substack frame ranges were adjusted (e.g., 101–2100-25, 301–2300-25) to capture periods of active migration. Cell tracking was performed via the *Manual Tracking*. Between 15–20 individual cells were tracked per AVI. Tracking data were saved as.txt files compatible with the Ibidi Chemotaxis and Migration Tool software.

### Chemotactic Analysis

Tracking files were imported into the Ibidi Chemotaxis and Migration Tool. Settings for analysis were as follows: Slices used: 1–80, X/Y calibration: 1.662◻µm per pixel, Time interval: 62.5◻s. X/Y calibration was determined from ZEN field-of-view data, with 851◻µm mapped across 512 pixels, yielding 1.662◻µm/pixel. All data were processed and interpreted according to Ibidi’s analysis guidelines. All tracking data are included in **Supplementary File_1** and **Supplementary File_2**, for *A. castellanii* and *A. polyphaga*, respectively.

We omitted the forward migration index (FMI), the projection of a trajectory onto the gradient axis divided by path length, ranging from −1 to +1, widely used to quantify chemotactic bias. In our μ-Slide setup, cells were seeded in the central channel, ADM was placed in the left reservoir, and chemoattractant in the right reservoir. Because the central channel is open to both reservoirs, the solute gradient relaxes continuously as the attractant diffuses across the channel and into the opposite reservoir. Without concurrent gradient tracing (e.g., a fluorescent tracer), the gradient direction and magnitude are time-varying, and may even invert late in the run. Since FMI assumes a fixed, known gradient axis, using it here would bias values toward zero (or spuriously negative) as the gradient flattens/changes, making FMI unreliable for inference. We therefore focused on gradient-agnostic metrics— velocity, directness (net/accumulated distance), and accumulated distance, benchmarked against ADM controls and supported by trajectory plots.

### Statistics

All quantitative data are reported as mean◻±◻standard deviation (SD), with each condition replicated in three biological replicates. Statistical differences between treatment groups and the ADM (control) were evaluated using Two-way ANOVA followed by Dunnett’s multiple comparisons test, which controls the family-wise error rate when comparing multiple treatments against a single control. A two-tailed *p*-value <0.05 was considered statistically significant.

## Results and Discussion

### Chemotactic responses of *A. castellanii* and *A. polyphaga* to microbial sugars and peptides

To revisit and expand upon early investigations into amoeboid chemotaxis (Schuster and Levandowsky, 1996) we evaluated the responses of *Acanthamoeba castellanii* and *A. polyphaga* to a panel of microbial surface-associated sugars (Mannan, Mannose, GlcNAc, and MurNAc), the bacterial peptide fMLP, and its analogue BOC-FLFLF. Using ibidi μ-Slide chemotaxis chambers and manual tracking across 2000 frames (2.5 s/frame), we analysed ∼40–50 cells per condition over three biological replicates. Tracking data were processed using the ibidi Chemotaxis & Migration Tool to generate trajectory maps and quantitative metrics of velocity, directness, and accumulated distance **(Figures 1–2)**.

**Figure 1.**
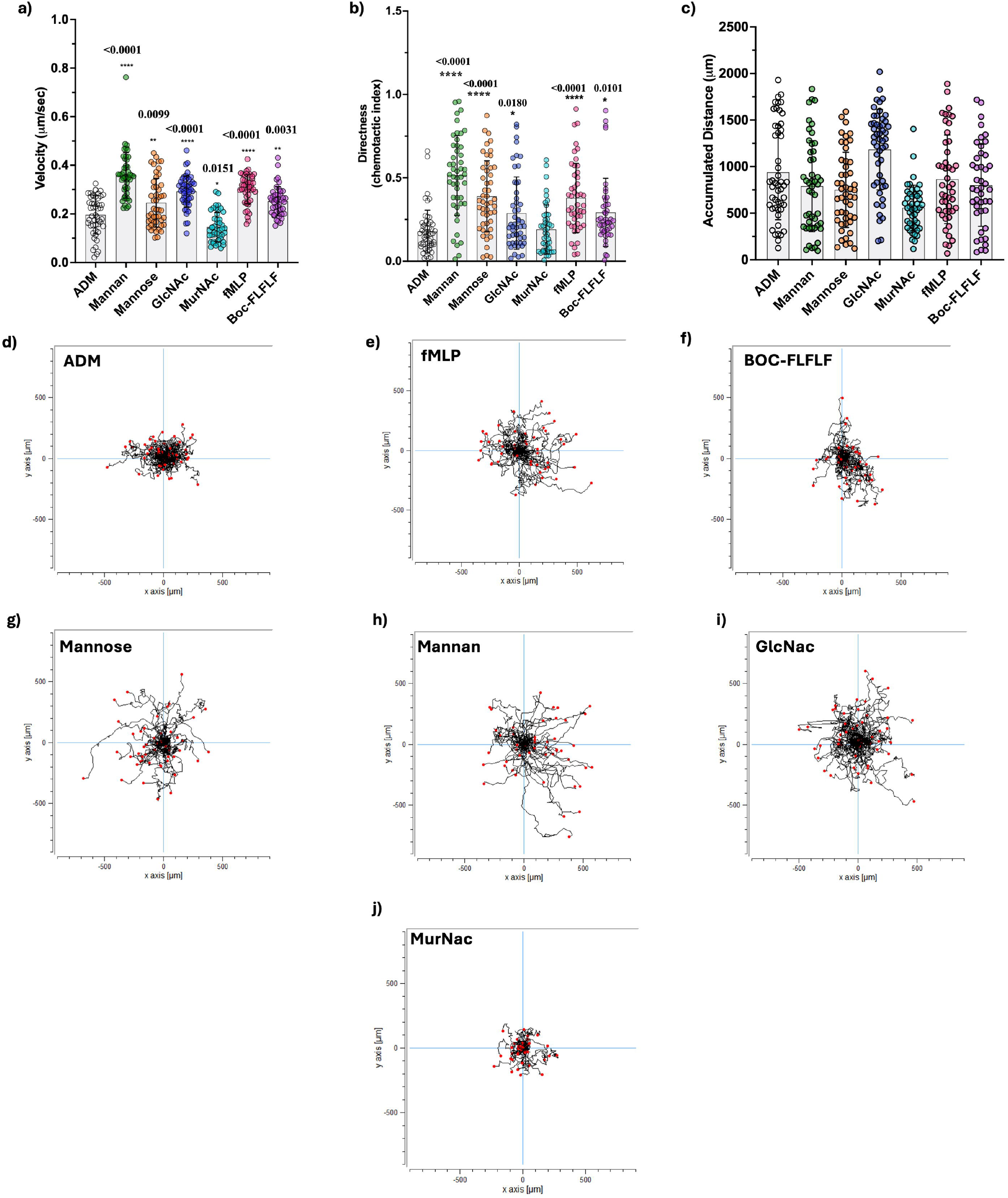
Chemotactic responses of *A. castellanii* to microbial ligands. Quantification of migration **a)** velocity, **b)** directness and **c)** accumulated distance in the presence of ADM (control), Mannan, Mannose, GlcNAc, MurNAc, fMLP, and BOC-FLFLF. Data represent means ± SD from three biological replicates. Asterisks indicate statistically significant differences relative to ADM (*p < 0.05, **p < 0.01, ****p < 0.0001).

**Figure 2.**
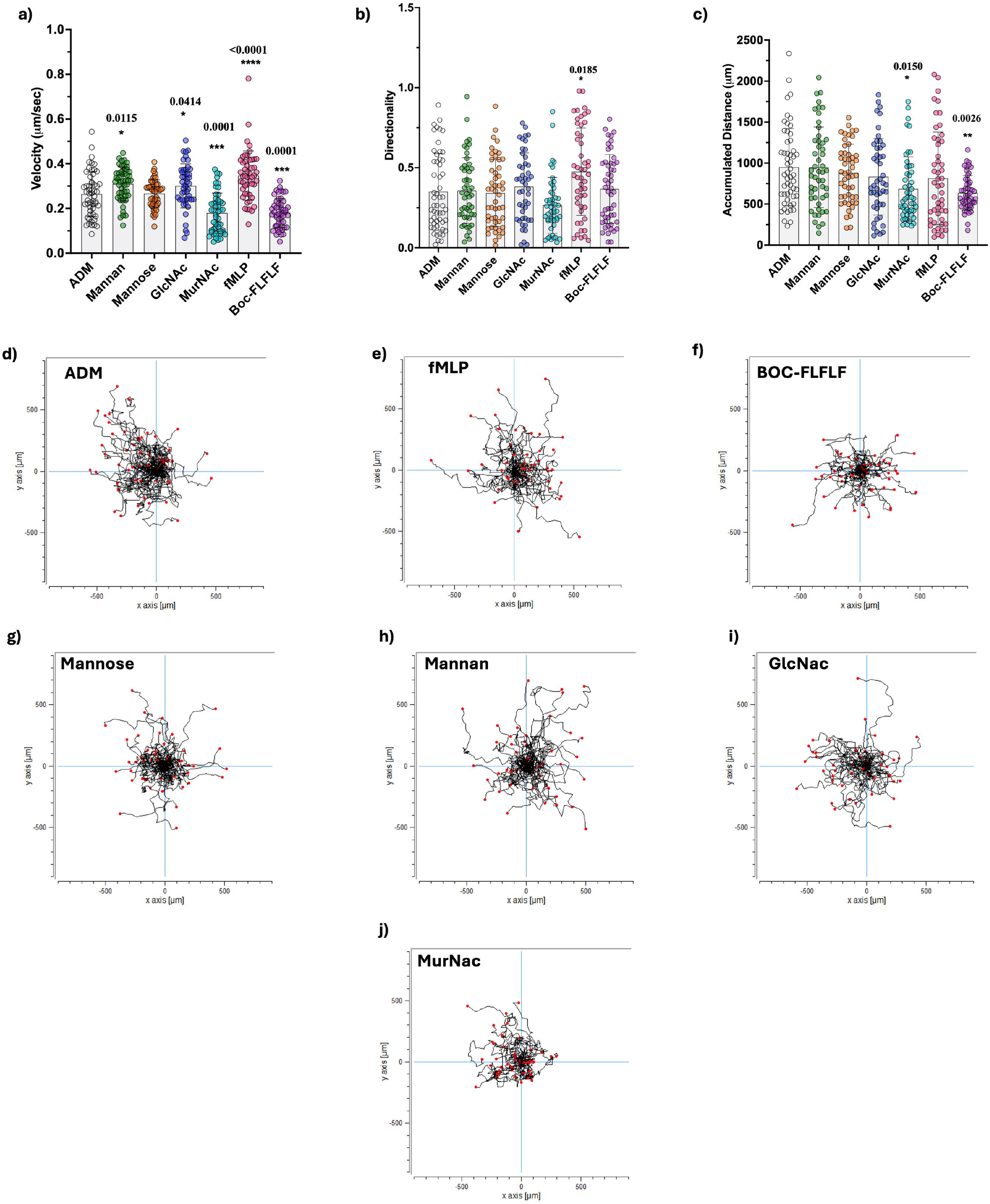
Chemotactic responses of *A. polyphaga* to microbial ligands. Quantification of migration **a)** velocity, **b)** directness and **c)** accumulated distance in the presence of ADM (control), Mannan, Mannose, GlcNAc, MurNAc, fMLP, and BOC-FLFLF. Data represent means ± SD from three biological replicates. Asterisks indicate statistically significant differences relative to ADM (*p < 0.05, **p < 0.01, ****p < 0.0001; Dunnett’s test).

Relative to the ADM baseline, *A. castellanii* showed significantly increased velocities in response to Mannan (p<0.0001), Mannose (p=0.0164), GlcNAc (p<0.0001) and fMLP (*p*<0.0001). Interestingly, BOC-FLFLF also produced a modest but significant increase (*p*<0.0059), while MurNAc showed statistically significantly lower velocity (*p*<0.0230). These findings highlight a preferential chemokinetic response toward mannose-rich glycans and bacterial peptides **(Figure 1a)**. Similarly, significant increases in directional persistence were observed for Mannan, Mannose, GlcNAc, fMLP, and BOC-FLFLF. Notably, mannan induced the highest directional bias (p < 0.0001). The stronger response to mannan relative to mannose is consistent with an avidity requirement, whereby multivalent mannose displays (e.g., mannan) cluster mannose-reactive lectins (e.g., MBP) and amplify downstream signalling, whereas monomeric mannose affords low-avidity engagement and weak directional bias (Casillo et al., 2021). At the nanomolar concentrations used, bulk viscosity is unlikely to account for the rightward bias; instead, mannan’s slower diffusion likely sustains a steeper, longer-lived gradient near the source, and limited surface adsorption could introduce a mild haptotactic component. Together with multivalent lectin engagement, these physical and biochemical factors plausibly underlie the stronger directional response relative to small sugars or fMLP.

MurNAc failed to enhance directionality significantly, suggesting these molecules may be insufficient cues for directional migration on their own **(Figure 1b)**. Unlike velocity and directness, accumulated distance did not significantly differ across conditions, suggesting that overall path length was not as sensitive an indicator of chemotactic response in this assay format. A borderline effect was noted with GlcNAc (*p*<0.0503), which may merit further exploration under longer timeframes or with automated tracking **(Figure 1c)**. Trajectory plots revealed pronounced directional migration toward fMLP, Mannan, and GlcNAc, while tracks for ADM appeared more isotropic. MurNAc and BOC-FLFLF produced reduced path lengths **(Figure 1d-j)**.

By contrast, *A. polyphaga* exhibited a more selective response profile **(Figure 2)**. Significantly increased velocities were observed only for Mannan (*p*<0.0115), GlcNAc (*p*<0.0414), and fMLP (*p*<0.0001), with fMLP producing the most pronounced effect. Directional persistence was significantly enhanced solely by fMLP ((*p*<0.0185), while other cues—including Mannose and GlcNAc—failed to elicit directional migration. Accumulated distance was significantly reduced in response to MurNAc (*p*<0.0150) and BOC-FLFLF (*p*<0.0026), suggesting inhibitory effects on sustained migration unique to this species (**Figure 2a–c)**. Trajectory plots show that *A. polyphaga* exhibits directional migration only toward fMLP, while responses to Mannan, Mannose, and GlcNAc appear non-directed, and MurNAc and BOC-FLFLF visibly suppress motility—highlighting a restricted chemotactic repertoire relative to *A. castellanii* **(Figure 2d-j)**.

Comparative analysis reveals that *A. castellanii* displays a broader and more robust chemotactic repertoire, responding to multiple glycans and peptides with increased speed and directional bias, whereas *A. polyphaga* shows a more constrained profile with directional responses largely limited to fMLP and reduced long-range migration in the presence of certain ligands. These patterns point to interspecies differences in receptor repertoires and signalling thresholds rather than a single shared pathway. For mannose-class cues specifically, the stronger response to mannan than mannose is consistent with multivalent engagement of a mannose-binding lectin; by contrast, responses to GlcNAc and to fMLP likely involve distinct lectin(s) and a peptide-sensing GPCR-like mechanism, respectively. Consistent with a multivalency model, pre-exposure of *A. castellanii* to free mannose yielded an “adhesion-without-uptake” phenotype—compatible with competitive masking of mannose-sensitive binding sites (e.g., MBP or related lectins) and insufficient multivalent engagement to trigger bacteria uptake **(Supplementary File. 3)**. This working model does not imply that MBP alone mediates uptake; parallel receptors and pathways may contribute. Nevertheless, this aligns with reports that soluble mannose analogues competitively mask mannose-sensitive lectins (Garate et al., 2005; Yoo and Jung, 2012).

Overall, *Acanthamoeba* chemotaxis appears modular, with ligand-specific pathways tuned differently across species; *A. castellanii*’s broader sensitivity may reflect a generalist foraging strategy, whereas *A. polyphaga*’s selective fMLP bias suggests tighter receptor gating or downstream signalling tuned to specific microbial cues. While both species remain valuable for comparative investigation, the broader chemotactic range of *A. castellanii* may provide a useful framework for probing general mechanisms of prey detection and microbe–protist interactions. Continued cross-species analyses will be essential to understand how recognition strategies vary across *Acanthamoeba* and what evolutionary or ecological factors drive this diversity. such as differences in MBP and lectin repertoires.

### Formyl peptide modulation differs between *Acanthamoeba* species

We sought to determine whether *Acanthamoeba* species exhibit mammalian-like formyl peptide modulation, particularly whether chemotactic responses to fMLP are antagonised by BOC-FLFLF, a known competitive inhibitor of the mammalian formyl peptide receptor (FPR). To test this, we exposed *A. castellanii* and *A. polyphaga* to fMLP alone, BOC-FLFLF alone, and a combination of both **(Figure 3)**.

**Figure 3.**
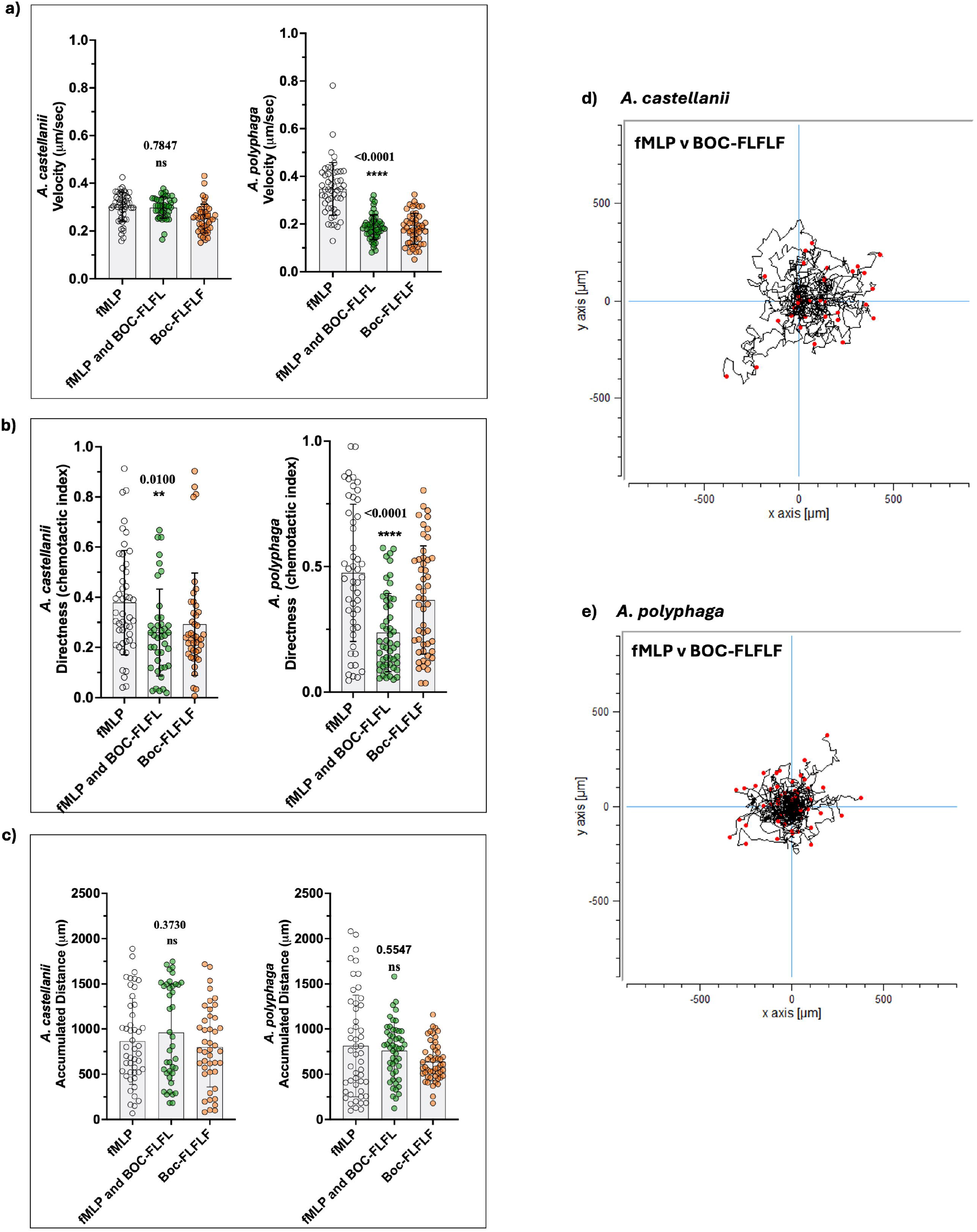
Effect of BOC-FLFLF on *A. castellanii* and *A. polyphaga* chemotaxis toward fMLP. Acanthamoeba responses were compared under three conditions: fMLP alone, fMLP + BOC-FLFLF and BOC-FLFLF alone. **a)** Migration velocity (µm/min) between fMLP and fMLP + BOC-FLFLF. **b)** Directness (chemotactic index) **c)** Accumulated distance travelled. Trajectory plot showing the effects of BOC-FLFLF on fMLP on **d)** *A. castellanii* and **e)** *A. polyphaga*. Data represent mean◻±◻SD from three biological replicates (n=≈45-55 cells/condition). Statistical comparisons were made using one-way ANOVA followed by Dunnett’s multiple comparisons test; significance levels: **\*\****p* < 0.01.

In neutrophils, fMLP is a potent chemotactic agonist, while BOC-FLFLF is a selective FPR antagonist that blocks fMLP-driven signalling (Johansson et al., 1993; Hayashi et al., 2014).

In *A. castellanii*, the addition of BOC-FLFLF to fMLP did not significantly (*p*>0.05) affect velocity or accumulated distance relative to fMLP alone, but both parameters remained comparable to BOC-FLFLF alone. However, directness was significantly reduced (*p*<0.01) compared to fMLP alone and matched the reduced directionality seen with BOC-FLFLF alone. These findings suggest that while BOC-FLFLF suppresses directional sensing in *A. castellanii*, it does not interfere with overall motility or path length. In contrast, *A. polyphaga* exhibited a broader suppressive response: the fMLP + BOC-FLFLF combination significantly reduced both velocity (*p*<0.0001) and directness (*p*<0.0001) relative to fMLP alone, while these reductions were not significant compared to BOC-FLFLF alone. Accumulated distance remained unchanged across conditions. This indicates that in *A. polyphaga*, BOC-FLFLF more broadly dampens chemotactic responses, reducing both speed and orientation, potentially reflecting lower signalling thresholds or less selective peptide sensing mechanisms **(Figure 3a-c)**. Trajectory plots further support these distinctions, revealing diminished and disoriented migration tracks in both species under co-treatment, but with more profound motility suppression in *A. polyphaga* **(Figure 3d-e)**.

These interspecies differences suggest that formyl peptide recognition and antagonism are modulated by divergent receptor sensitivities or signalling architectures. The ability of *A. castellanii* to sustain motility while selectively losing directional bias may indicate the presence of a more finely tuned sensing system. These behavioural outcomes, particularly the selective suppression of directness in *A. castellanii* without impairment of motility, are reminiscent of GPCR-regulated chemotactic adaptation observed in neutrophils and *Dictyostelium* systems. In human neutrophils, CAPRI (a calcium-promoted Ras inactivator) is essential to regulate GPCR-mediated Ras activation and adaptation, thereby controlling cell sensitivity across a wide range of chemoattractant concentrations. CAPRI also fine-tunes directional sensing by locally deactivating Ras following fMLP stimulation (Xu et al., 2021a; Xu et al., 2021b). The broader suppression of both velocity and directionality in *A. polyphaga* upon co-treatment with BOC-FLFLF may reflect species-specific differences in receptor signalling strength, ligand binding affinity, or the responsiveness of downstream Ras regulatory networks. Although canonical FPR homologs remain unidentified in *Acanthamoeba*, the functional parallels we observe, particularly the selective antagonism of directional sensing, strongly suggest the existence of a divergent yet functionally analogous GPCR–Ras signalling circuit mediating peptide-driven chemotaxis. Indeed, genomic analysis reveals that *A. castellanii* encodes at least 35 putative GPCRs across several classes (families 1, 2, and 6) and five G-protein α-subunit genes, implying a sophisticated signalling network capable of discriminating peptide-based cues (Clarke et al., 2013). Genomic context supports a mechanistic basis for divergence: *A. castellanii* (Neff) harbours multiple GPCR families and lectins in a curated genome, whereas *A. polyphaga* currently has a larger, draft nuclear assembly with less complete receptor annotation. Together with species-specific microbial associations (e.g., giant virus interactions in *A. polyphaga*) (Chelkha et al., 2018), these differences plausibly tune receptor repertoires and signalling thresholds, aligning with the broader *A. castellanii* and more selective *A. polyphaga* chemotactic signatures we observed.

## Conclusion

By revisiting and expanding upon the foundational chemotaxis work by Schuster and Levandowsky (Schuster and Levandowsky, 1996) with micro-fluidic gradients and frame-by-frame tracking, we validate most of this classic study. *A. castellanii* migrates strongly to Mannan, fMLP and GlcNAc and shows little response to MurNAc or BOC-FLFLF but revealed three new layers of insight. First, stable nano-litre gradients uncovered clear attraction to mannan/mannose that the earlier millimetre-scale slide could not resolve, consistent with the species’ mannose-binding–proteins (Corsaro, 2022). Second, comparative analysis reveals that *A. castellanii* exhibits a broader and more nuanced chemotactic repertoire than *A. polyphaga*, with stronger responses to multiple glycans and peptides with a more robust directional sensing. In contrast to *A. polyphaga* which displayed a narrower chemotactic profile, characterized by selective sensitivity to fMLP and broader suppression in response to BOC-FLFLF. Third, these behavioural contrasts are compatible with GPCR-Ras adaptation paradigms borrowed from neutrophils and *Dictyostelium*, where Ras-GAP tuning can uncouple velocity from directional bias (Duda-Chodak et al., 2015), although the underlying receptors in Acanthamoeba remain unidentified.

Although both species are frequently isolated from the same environmental sources, including soil, freshwater systems, drinking water, air, and biofilms, with no published evidence supporting distinct niche partitioning, we hypothesise that their divergent chemotactic “signatures” reflect micro-niche specialisation, fine-tuning of receptor repertoires to local prey spectra rather than habitat partitioning. Our data establish quantitative benchmarks that can guide subsequent efforts to link ligand classes to specific Acanthamoeba receptors, pending targeted genetic and pharmacological validation.

## Abbreviations

fMLP: *N*-formyl-methionyl-leucyl-phenylalanine (bacterial formyl peptide chemoattractant)
BOC-FLFLF: tert-butyloxycarbonyl-Phe-Leu-Phe-Leu-Phe (formyl peptide receptor antagonist)
ADM: Acanthamoeba Defined Media, neutral control medium used in chemotaxis assays
GPCR: G-protein coupled receptor
MAMP: Microbe-associated molecular pattern
MBP: Mannose-binding lectin or protein
GlcNAc: *N*-acetyl-d-glucosamine (a bacterial sugar ligand)
MurNAc: *N*-acetyl-muramic acid (peptidoglycan component)
PYG: Peptone-Yeast-Glucose medium (for growing *Acanthamoeba*)
MBP1 / LBP: Mannose-binding protein 1 or Laminin-binding protein

## Data availability

All data generated or analysed during this study are included in the manuscript and supporting files.

## Acknowledgements

F.N. conceptualized and designed the study; V.H and F.N performed the experiments and data analysis, F.N. drafted the manuscript, V.H, BW.W and F.N edited the manuscript.

