## Supplementary File 3 for "Re-evaluating of Chemotaxis in *Acanthamoeba castellanii* and *Acanthamoeba polyphaga*: A Modern Perspective Using Advanced Imaging and Tracking Technologies"

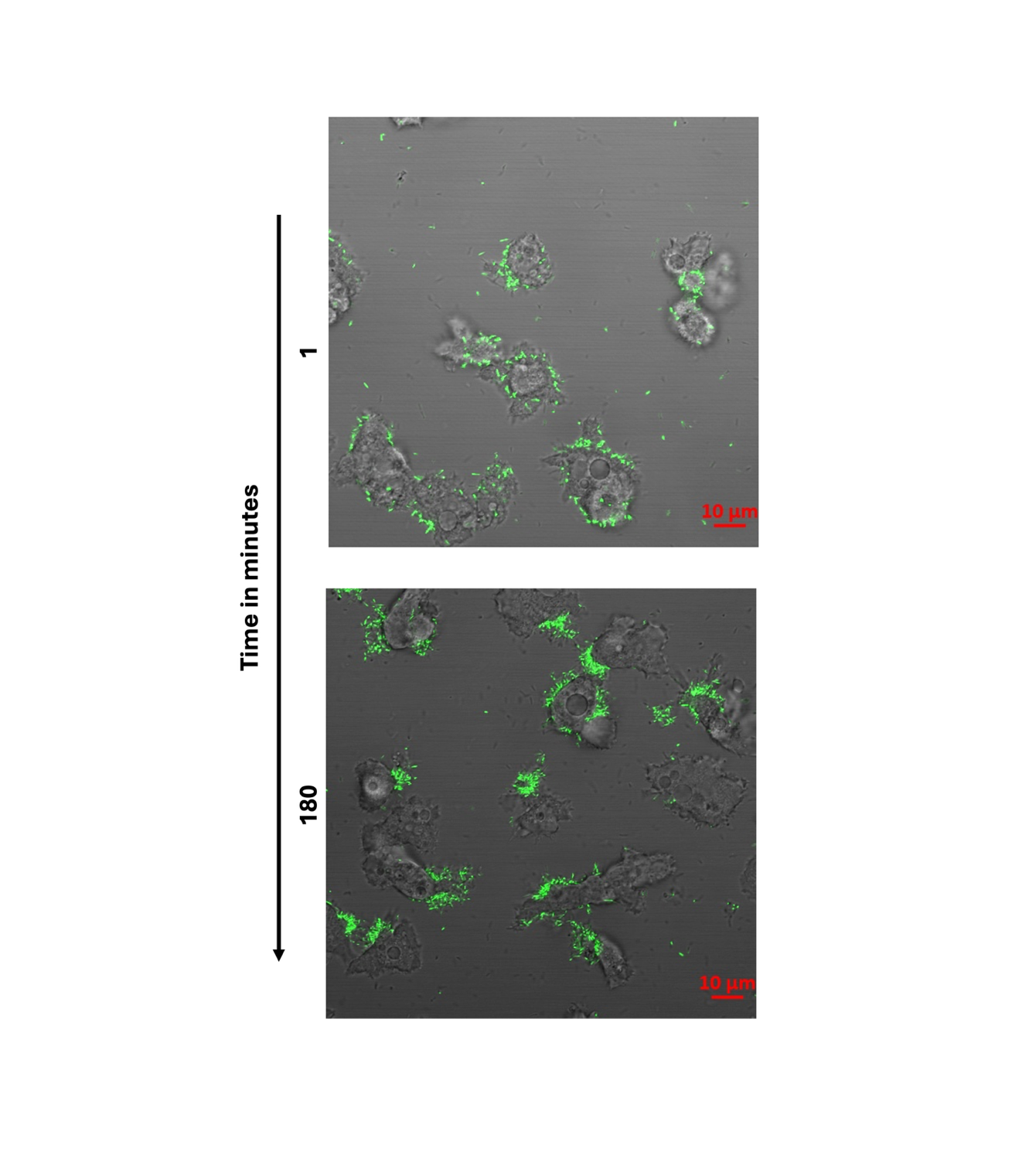

**Supplementary Figure 1: Mannose pre-exposure promotes accumulation of bacteria on the surface Acanthamoeba castellanii.** Trophozoites were pre-exposed to 1 mM of **D-mannose** for 30 min in ADM, then infected with **GFP-expressing** C. jejuni at **MOI = 100**. Interactions were imaged live in ADM for ~3 h. Following mannose pre-exposure, bacteria accumulate along the amoeba surface without internalization over time, consistent with **adhesion-without-uptake** under these conditions. **Zeiss LSM880** Differential interference contrast (DIC) (grey), overlaid with GFP fluorescence (green). Representative fields are shown at **t = 1 min** (top) and **t = 180 min** (bottom). **Scale bar: 10 µm.**
